# Adaptive plasticity in the healthy reading network investigated through combined neurostimulation and neuroimaging

**DOI:** 10.1101/2022.04.20.488885

**Authors:** S. Turker, P. Kuhnke, F. R. Schmid, V. K. M. Cheung, B. Zeidler, K. Seidel, L. Eckert, G. Hartwigsen

## Abstract

The reading network in the human brain comprises several regions, including the left inferior frontal cortex (IFC), ventral occipito-temporal cortex (vOTC) and dorsal temporo-parietal cortex (TPC). The left TPC is crucial for phonological decoding, i.e., for learning and retaining sound-letter mappings. Here, we tested the causal contribution of this area for reading with repetitive transcranial magnetic stimulation (rTMS) and explored the response of the reading network using functional magnetic resonance imaging (fMRI). 28 healthy adult readers overtly read simple and complex words and pseudowords during fMRI after effective or sham TMS over the left TPC. Behaviorally, effective stimulation slowed pseudoword reading. A multivariate pattern analysis showed a shift in activity patterns in the left IFC for pseudoword reading after effective relative to sham TMS. Furthermore, active TMS led to increased effective connectivity from the left vOTC to the left TPC, specifically for pseudoword processing. The observed changes in task-related activity and connectivity suggest compensatory reorganization in the reading network following TMS-induced disruption of the left TPC. Our findings provide first evidence for a causal role of the left TPC for overt pseudoword reading and emphasize the relevance of functional interactions in the healthy reading network for successful pseudoword processing.

## 1. Introduction

Reading is a core feature of human communication and crucial for participating in everyday social life, work and interpersonal communication. Fluent reading is based upon multiple, hierarchically organized processes, including orthographic recognition, orthographic-phonological mapping (i.e., decoding), and semantic access (Xia et al., 2017). Known words, regardless their complexity, are usually automatically accessed as whole word forms in the mental lexicon (so-called sight word reading) (Ehri, 2005). Reading this sentence took you probably only a few seconds although you had to process the pronunciation and meaning of eighteen words consecutively and at a high speed. The single sounds that make up these words were most likely beyond your awareness, i.e., you did not have to look at single letters or decode them for reading the sentence. When reading pseudowords (i.e., words without any meaning), on the other hand, you rely considerably on the single sounds and/or the syllables they occur in. Therefore, reading (complex) pseudowords is more challenging for typical adult readers because it is a non-automatic reading process.

The universal reading network in the human brain supporting these processes comprises three major circuits: (i) the left inferior frontal cortex (IFC), (ii) the left dorsal temporo-parietal cortex (TPC) and (iii) the left ventral occipito-temporal cortex (vOTC) (Pugh et al., 2001; Rueckl et al., 2015). Neuroimaging studies suggest that the left IFC is involved in various processes, including attention and language functions (e.g., phonological output resolution; Taylor et al., 2013). The left TPC, often referred to as the ‘decoding’ center in the human brain, is responsible for the transformation of orthographic elements into associated phonological codes (Linkersdörfer et al., 2012). The last region, the left vOTC, shows growing sensitivity to print during reading acquisition (Chyl et al., 2021) and optimizes linguistic processing for quick access to familiar words by filtering out meaningless grapheme strings (e.g., pseudowords) (Gagl et al., 2020).

Although non-invasive brain stimulation (NIBS) studies have proliferated over the past two decades, only few studies have applied NIBS to specifically modulate reading-related processes, and practically none have explored the effect of NIBS on overt pseudoword reading (Turker & Hartwigsen, 2021a). Understanding the causal engagement and functional contribution of brain areas to reading, however, is vital to improve our understanding of the healthy reading network. Additionally, it can help uncover and alleviate potential deficits encountered by individuals with dyslexia (see Turker & Hartwigsen, 2021b). Existing NIBS studies with typical readers provide first evidence for causal roles of reading-related regions to different reading subprocesses (see summary in Turker & Hartwigsen, 2021a). For instance, the left anterior and posterior IFC contribute to phonological and semantic aspects of reading, respectively (e.g., Devlin et al., 2003; Gough et al., 2005; Hartwigsen et al., 2010 a,b). Other studies further highlight the left vOTC as critical region for word processing (Duncan et al., 2010) but also confirm its relevance for pseudoword processing (Pattamadilok et al., 2015). Finally, the left TPC was identified as key area for phonological processes related to reading, in line with its expected role as grapheme-phoneme-conversion center (e.g., Costanzo et al., 2012; Liederman et al., 2003). However, existing NIBS studies on pseudoword reading focused on relatively simple bisyllabic pseudowords and included few trials. Moreover, none of the previous studies used neuroimaging to explore the underlying neuronal changes induced by neurostimulation. Consequently, the underlying neural correlates of stimulation-induced modulation of the reading network remain unclear.

Likewise, NIBS studies with atypical readers support the causal role of the left TPC to reading, but neglect underlying neural changes and mechanisms (see review by Turker & Hartwigsen, 2021b). Behaviorally, several single- or multiple-session studies with individuals with dyslexia found NIBS-induced improvements in reading performance in low frequency word, pseudoword and text reading after facilitation of the left TPC, or simultaneous inhibition of the right TPC and facilitation of the left TPC (e.g., Costanzo et al., 2016, 2019; Lazzaro et al., 2020, 2021). This further supports the critical role of the left TPC for reading processing but leaves open questions regarding potential NIBS-induced changes on the neural level, such as the potential of NIBS to modify the reading network transiently or permanently.

What are the neural correlates of the observed NIBS-induced behavioral modulation of reading performance in typical and atypical readers? On the one hand, facilitatory and inhibitory NIBS most likely either in- or decrease functional brain activation in the targeted area, which should in turn map onto changes in behavioral performance (Miniussi et al., 2013). Since individuals with dyslexia across all age groups show less engagement of the left TPC during reading when compared to typical readers (Richlan et al., 2011, 2013; Turker, 2018), it seems likely that inhibitory NIBS should result in a deterioration of reading performance in typical readers, whereas facilitation should help improve reading skills in atypical readers. On the other hand, impairing a core node of a network most likely results in significant up- and down-regulations in tightly connected brain regions, both in terms of functional activation and connectivity (Sale et al., 2015). Such potential compensatory mechanisms should occur after disruption of the targeted area, as described in previous combined TMS-fMRI studies in the language network and may be correlated with changes in behavioral performance (see Hartwigsen et al., 2013; Hartwigsen et al., 2017). Indeed, neuroimaging suggests that phonological deficits in children with dyslexia could stem from a disruption of functional connectivity between the three core reading areas (left IFC, vOTC and TPC) (van der Mark et al., 2011; Schurz et al., 2015), highlighting the importance of considering within-network-interaction. In terms of reading, it remains yet to be explored whether NIBS can alter functional activation and connectivity within the reading network, and whether this directly maps onto reading performance.

In the present study, we tested: (i) the causal contribution of the left TPC for efficient word and pseudoword reading by inducing a focal perturbation with repetitive transcranial magnetic stimulation (rTMS); and (ii) explore the reading network’s response to perturbation in terms of functional activation and connectivity. We hypothesized that the inhibition of the left TPC would lead to an increase in reading times and a decrease in accuracy for simple and complex pseudowords, with a stronger effect on complex pseudowords. Our second hypothesis was that the disruption of the left TPC should lead to an up-regulation of the left IFC and vOTC (i.e., higher activation within these areas and higher functional coupling with the disrupted region), and potentially also the contralateral right TPC (see Hartwigsen & Volz, 2021). This would be in line with our hypotheses that a disruption of the reading circuit requires functional reorganization and compensatory mechanisms to sustain reading. To test these hypotheses, we applied offline effective or sham rTMS to the left TPC of healthy adults who then performed a reading task during functional MRI. Subjects were asked to read aloud easy and complex words and pseudowords (see **Figure 1**). As main findings, we observed that effective rTMS (relative to sham stimulation) led to (1) slower speech onsets for pseudowords, (2) a shift in task-related activation patterns in the left IFC during pseudoword reading, and (3) stronger task-related functional coupling between the left vOTC and the left TPC.

**Figure 1.**
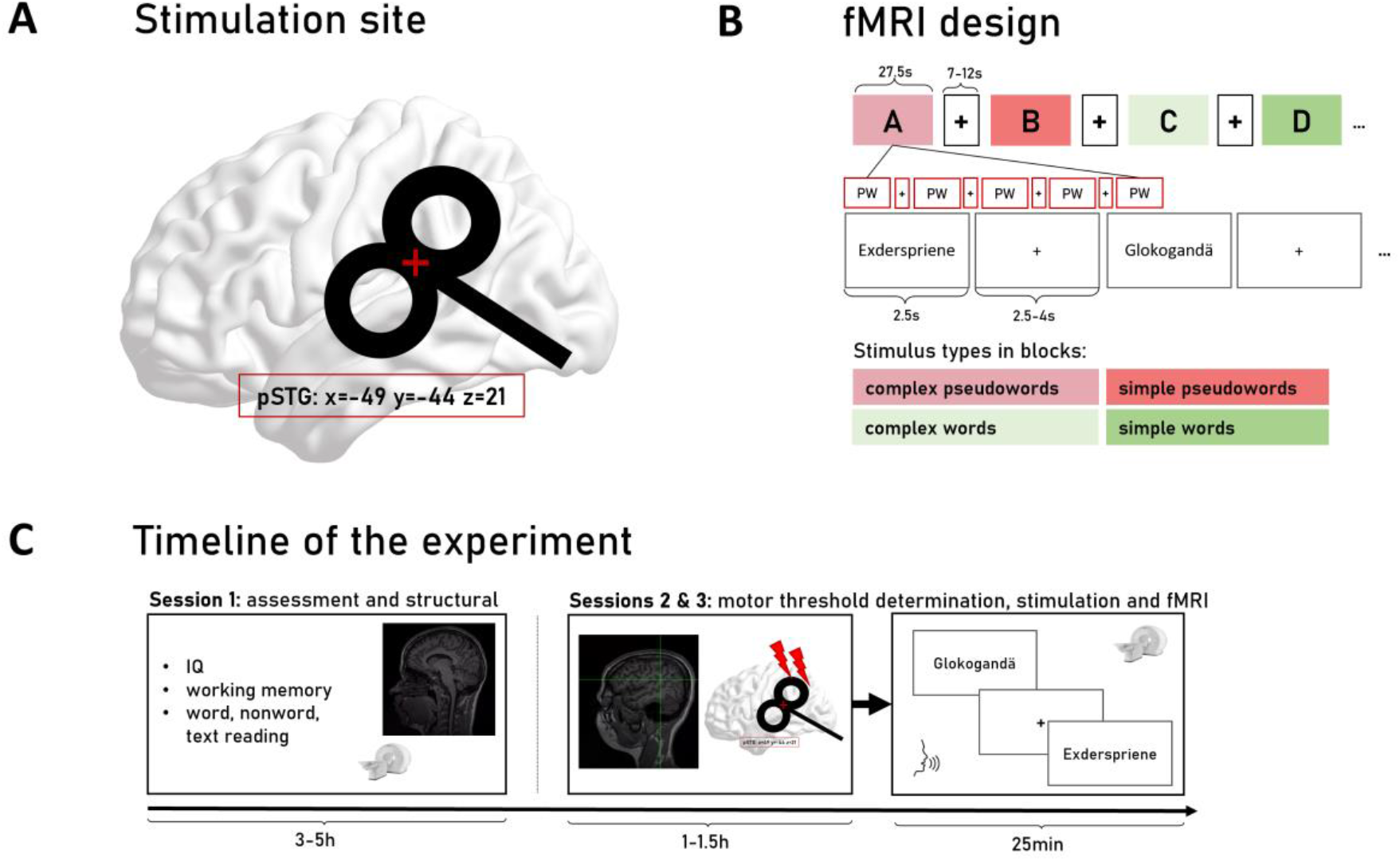
Details on stimulation site (A), fMRI design (B) and timeline of the experiment (C). The target was chosen based on previous meta-analyses and was situated in the left pSTG (−49/-44/21). During fMRI, subjects read simple and complex words and pseudowords (200 trials per scanning session; total duration: ∼ 25 minutes).

## 2. Results

### 2.1. Behavioral findings

We modelled the speech onset and accuracy of each trial using generalized linear mixed models (Tables 1-3). As fixed effects, the interaction between TMS (effective vs. sham), complexity (simple vs. complex stimuli), and stimulus type (words vs. pseudowords) and lower order terms were included in the model. A maximal random effects structure was used for each model to guard against inflated Type I errors. Results for speech onsets are displayed in **Tables 1 and 2** and visualized in **Figure 2**, results for reading accuracy are provided in **Table 3**.

**Table 1.** Effects of TMS on speech onsets. Speech onsets of each trial were modelled to follow a Gamma distribution using a generalized linear mixed model. Significance of predictors were assessed using the Wald test. *: p < 0.05, **: p < 0.01, ***: p < 0.001

| Predictor | $\chi^2(1)$ | p-value |
| --- | --- | --- |
| Intercept | 1614.6620 | $< 2.20 \times 10^{-16}$ *** |
| TMS | 0.0039 | 0.950 |
| Complexity | 64.2319 | $1.11 \times 10^{-15}$ *** |
| **Pseudoword** | 148.9187 | $< 2.20 \times 10^{-16}$ *** |
| **TMS x Complexity** | 1.9935 | 0.158 |
| **TMS x Pseudoword** | 3.7938 | 0.051 |
| **Complexity x Pseudoword** | 42.2177 | $8.17 \times 10^{-11}$ *** |
| **TMS x Complexity x Pseudoword** | 0.8281 | 0.363 |

**Table 2.**
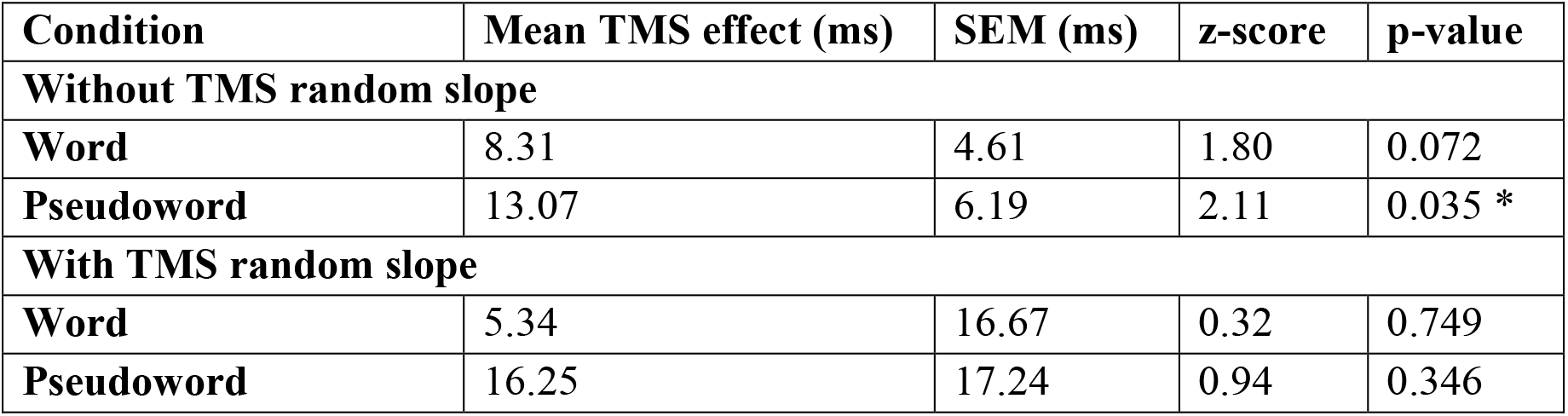
Marginal effect of TMS (effective – sham) on speech onsets for pseudowords using step-down analysis. Models with random subject intercepts and with or without random slopes for TMS were fitted. Significance predictors were assessed using the Wald test. *: p < 0.05, **: p < 0.01, ***: p < 0.001

**Table 3.**
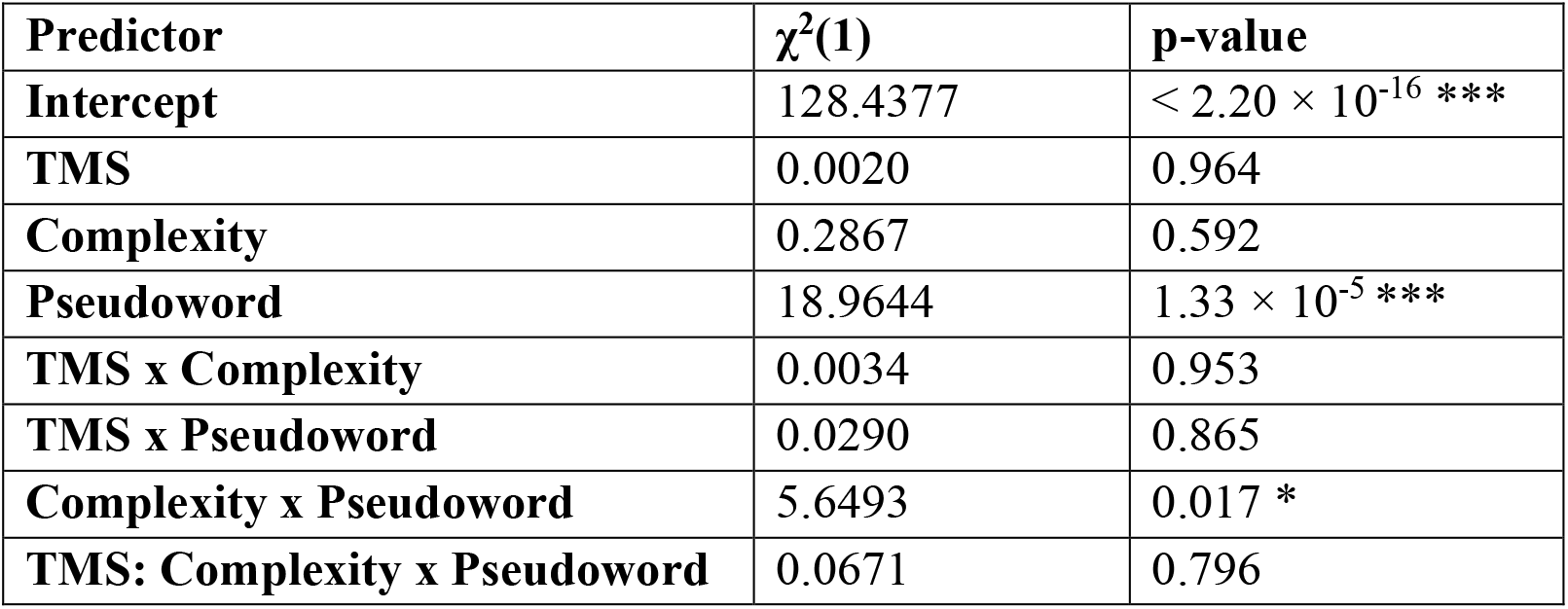
Effects of TMS on response accuracy. Correct and incorrect responses of each trial were modelled using a binomial generalized linear mixed model. Significance predictors were assessed using the Wald test. *: p < 0.05, **: p < 0.01, ***: p < 0.001

**Figure 2.**
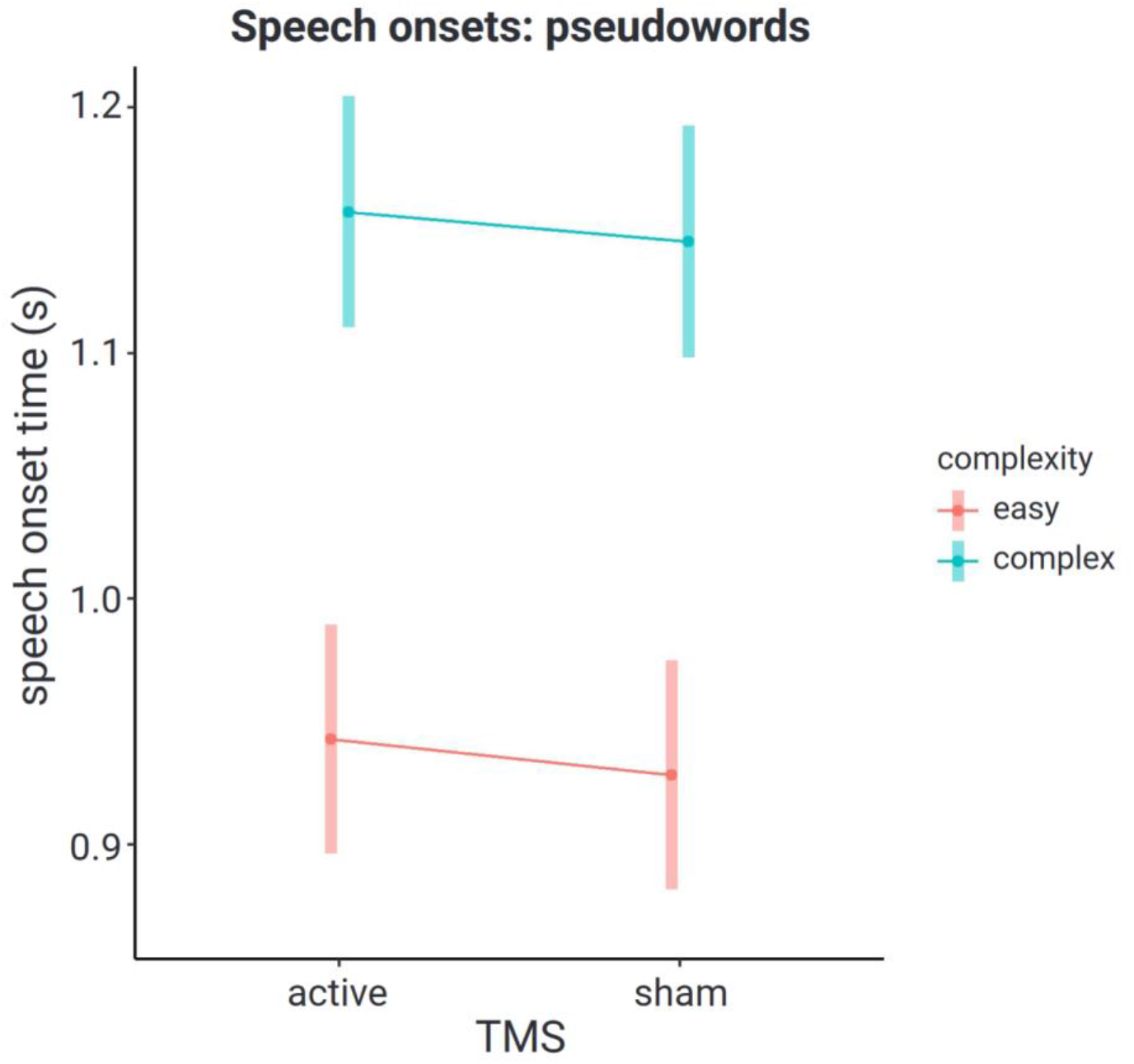
TMS effect on pseudoword speech onsets. (active stimulation vs. sham stimulation) modelled by word complexity (0 = simple; 1= complex)

The model shows that participants generally showed significantly longer speech onset times for pseudowords compared to words and for complex compared to simple stimuli. Likewise, subjects showed lower accuracy for pseudoword as compared to word reading. Significant complexity-by-pseudoword interactions furthermore indicated that these effects were enhanced for complex pseudowords. As for TMS effects, we found a marginally significant TMS-by-pseudoword interaction (p=0.051) for speech onsets (**Table 1**). Step-down analyses (**Table 2**) suggest that the effect was driven by a delay of 13 – 16 ms in pseudoword reading. Interestingly, the TMS effect on pseudowords was significant when the model included random intercepts (p=0.035), but not anymore when we included randoms slopes. This suggests large inter-individual variability in TMS response, which makes effects that generalize to the population hard to detect. We did not find any effect of TMS on reading accuracy (**Table 3**).

### 2.2. Functional activation

To explore differences in reading processing for stimulus type and complexity, we performed within-session univariate whole-brain analyses of the sham condition (see **Figure 3**). Pseudowords (bi- and four-syllabic) activated the left motor cortex, the left IFC, the bilateral posterior parietal cortices and the bilateral vOTC more than words. While activation of the motor cortex is most likely tied to higher articulation demands, the engagement of the other areas seems to be related to higher reading demands as compared to word processing. Words, on the other hand, led to higher activation in brain areas corresponding to the default mode network (DMN) (Smallwood et al., 2021). These include the bilateral angular gyri, middle temporal gyri, middle frontal cortices, and bilateral medial brain regions including the medial prefrontal cortices and the posteromedial cortices. Since sight word reading is an automatized, higher-order cognitive task, it is little surprising that word reading recruits areas of the DMN more strongly than pseudoword reading. Especially since word reading is reliant upon lexical retrieval and semantic access, which are known to depend on processing in the left angular gyrus and the left middle temporal cortex to left anterior temporal lobe. Recently, the role of the posterior parietal cortex for reading was confirmed in a few NIBS studies (Turker & Hartwigsen, 2021a for review). A stronger engagement of the visual word form area and its homologue for pseudoword processing, on the other hand, have not been explicitly discussed in research to date. Regarding complexity, we find primarily higher activation for complex stimuli in the bilateral motor cortices, the bilateral superior temporal gyri and areas within the posterior parietal cortex and occipital lobes.

**Figure 3.**
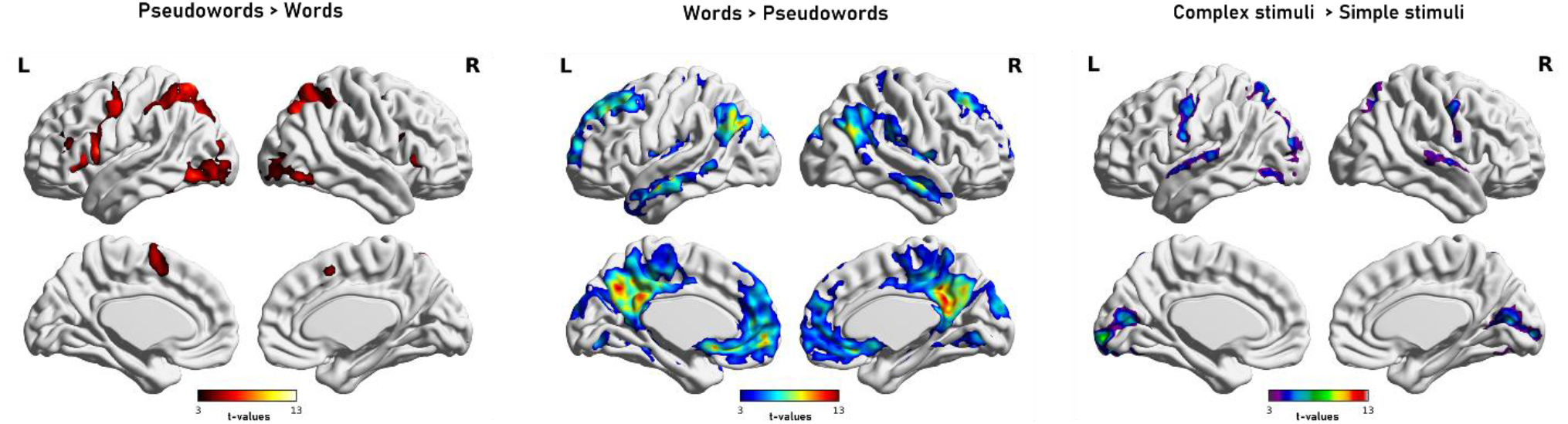
Univariate contrast maps. Results show functional brain activation differences due to stimulus type (pseudowords vs. words) and complexity ((complex pseudowords + complex words) > (simple pseudowords + simple words)) corrected at p<0.001 voxel-level and FWE-cluster-corrected at p<0.05.

Regarding TMS effects, univariate whole-brain analyses showed no significant differences in brain activation when comparing effective and sham TMS, not even for complex pseudowords that have the highest decoding demand. To better understand this, we performed an exploratory subject-specific analysis with individual subject maps thresholded at p<.001 (uncorrected) (**Figure 4**). Supporting our observation of large inter-individual variability in behavioral response to TMS, participants also showed large differences in univariate brain activation involving up- and down-regulation of the bilateral motor cortices and the occipital lobe bilaterally following effective TMS. It is particularly striking that the increased and decreased activation in response to effective TMS comprises largely overlapping areas in these regions. In other words, some individuals responded with more and others with less activation in the bilateral motor areas, portions of the superior temporal gyrus, and occipital areas in response to effective TMS as compared to sham TMS. These preliminary analyses show that neural responses to TMS differ considerably between individuals, which might explain the lack of group-level differences.

**Figure 4.**
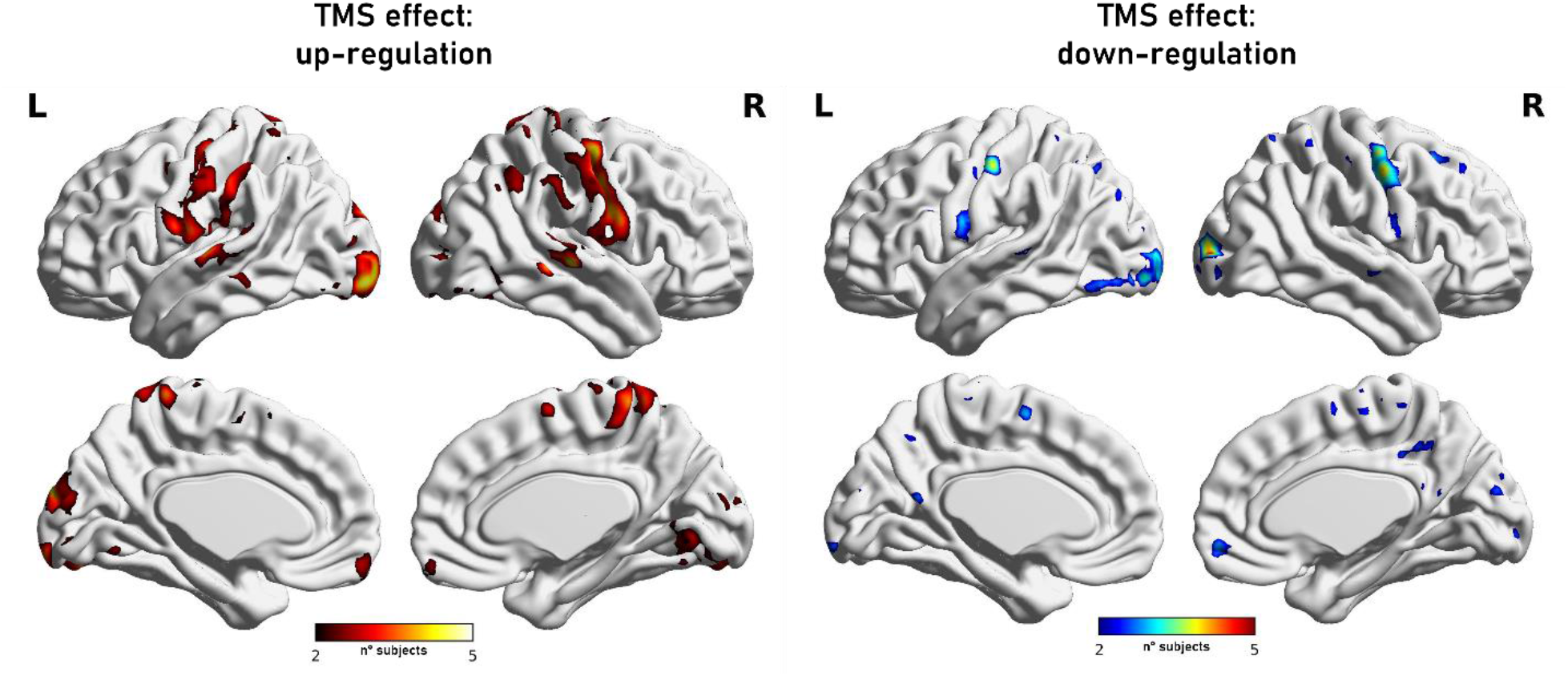
Exploratory individual subject maps of functional activation in response to effective TMS as compared to sham. Results on the left side show an up-regulation, i.e., a higher activation in brain areas after effective stimulation. Results on the right side show areas that responded to effective TMS with a down-regulation, i.e., with less brain activation. (p<0.001 uncorrected at the voxel level thresholded at 2 subjects; colour indicates number of subjects showing increase/decrease in the same voxel).

Since univariate analyses are often claimed to be insensitive to fine-grained differences in multi-voxel activity patterns we additionally performed a multivariate pattern analysis (MVPA) to investigate the effect of TMS on functional activation patterns within the core reading areas. Specifically, we used a spherical 5-mm “searchlight” across the whole brain, and at each searchlight location, we trained a machine learning classifier to decode between effective and sham TMS across participants, separately for words and pseudowords. We found above-chance between-subject decoding in the left posterior IFC selectively for pseudowords (**Figure 5**) (see a discussion on the role of the left mid to posterior IFC for pseudoword processing in Turker & Hartwigsen, 2021b). No effects were observed for word processing, supporting our hypothesis that effective stimulation of the left TPC primarily influences pseudoword processing. These findings indicate that effective TMS over the left TPC altered fine-grained multi-voxel activity patterns for pseudoword reading in the left IFG (pars triangularis) by resulting in a shift of the task-related activity pattern in that area.

**Figure 5.**
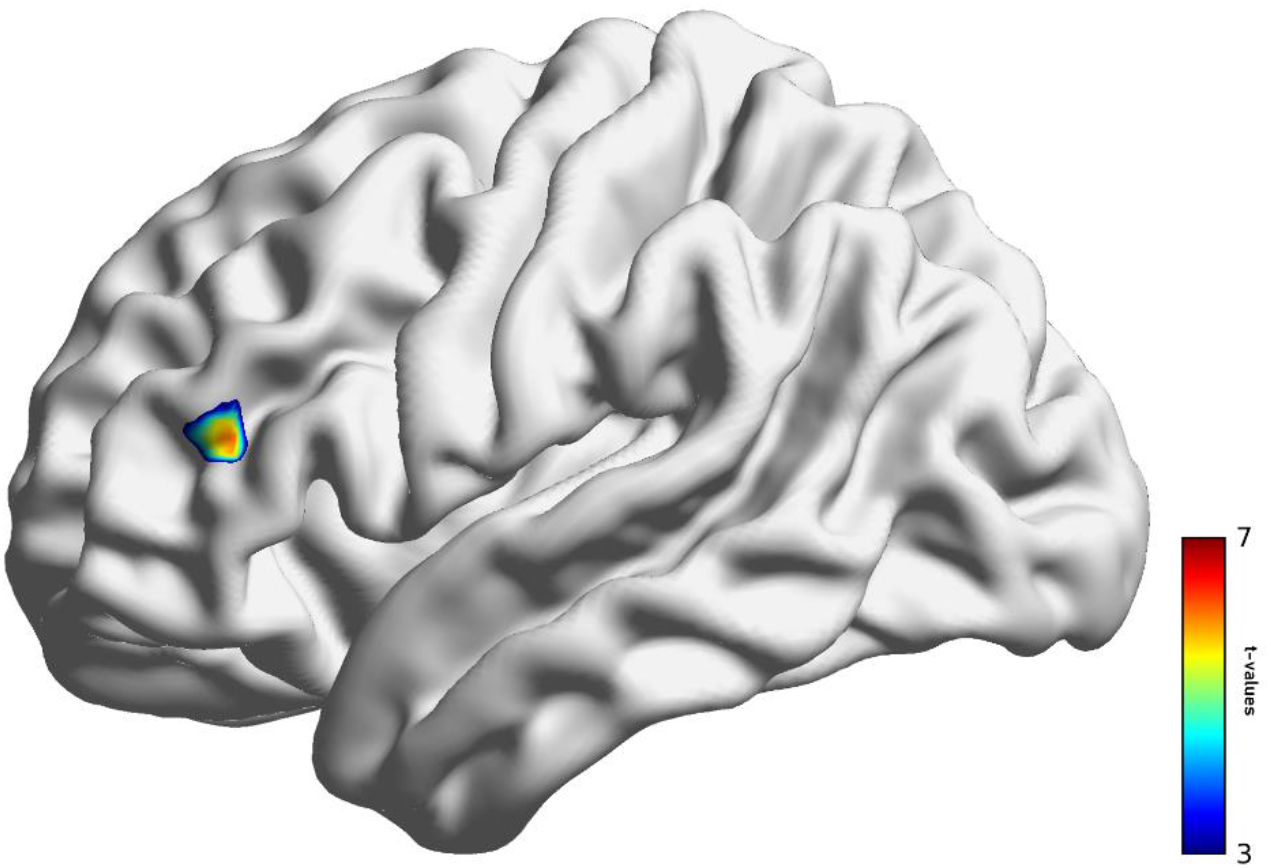
Results for between-subject searchlight MVPA. Following effective stimulation of the left TPC as compared to sham stimulation, we find a shift in functional activation patterns in the left IFC selectively for pseudoword reading. Activity patterns comprised beta estimates for each mini-block (see **Figure 1**) of every participant.

### 2.3. Effective connectivity

Finally, we performed Dynamic Causal Modeling (DCM) to map TMS-induced changes in effective connectivity within the reading network. The DCM model included the core reading areas (the left IFC, the left vOTC and the left TPC) that represent core nodes of the reading network that were also active for pseudoword and word reading, independent of TMS. We performed a DCM group analysis using Bayesian Model Reduction (Friston et al., 2016; Zeidman et al., 2019 a,b). To this end, we first defined a ‘full’ DCM model for each subject. According to this model, all three areas were bidirectionally connected, words and pseudowords could serve as driving inputs to every region, and each between-region connection could be modulated by effective and sham TMS. Bayesian Model Reduction then compared this full model to numerous reduced models. Finally, we computed the Bayesian model average, which is the average of parameter values across models weighted by each model’s probability, and thresholded the BMA at 99% parameter probability (see **Figure 6**).

**Figure 6.**
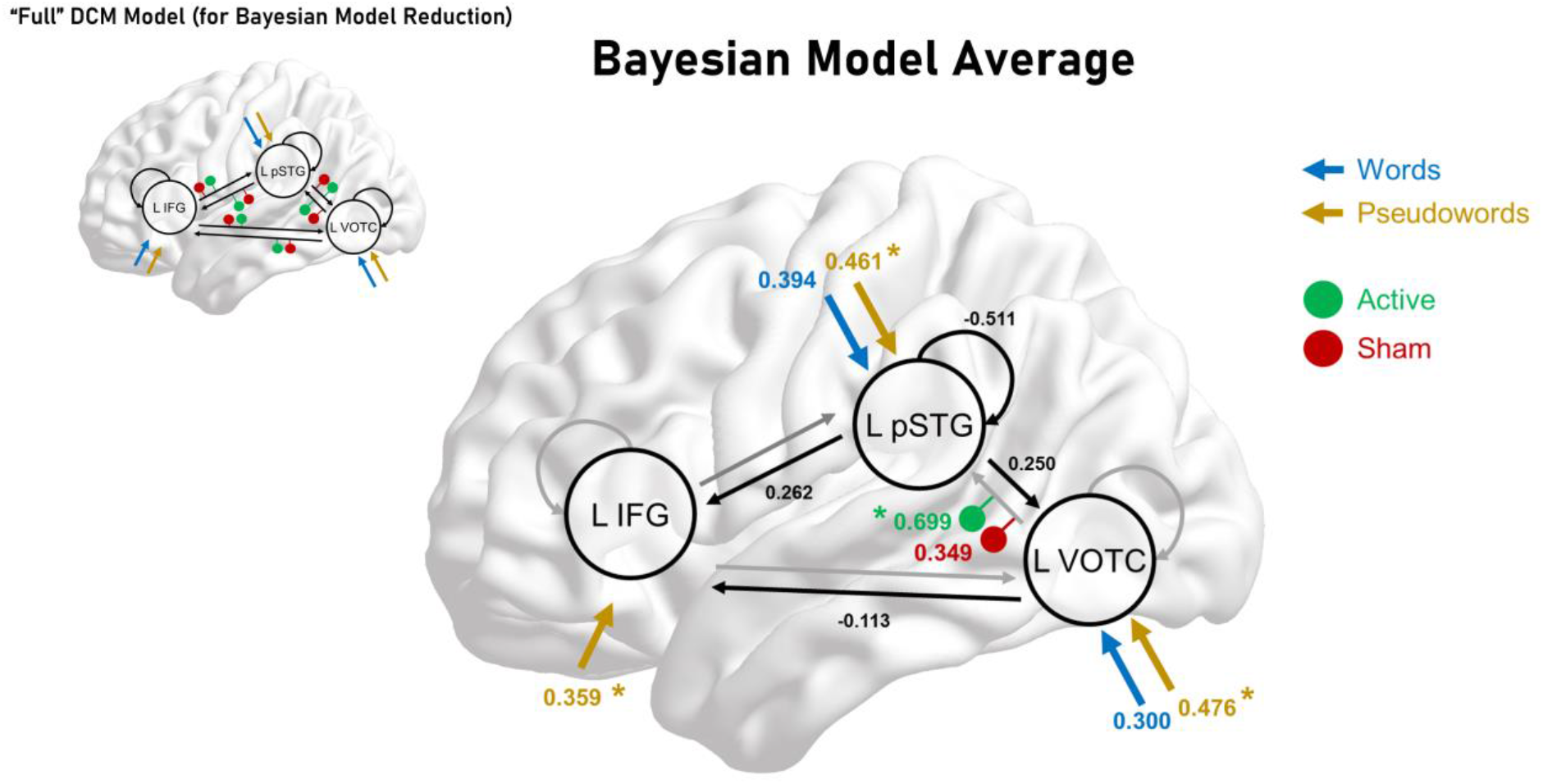
Dynamic causal modeling results. *Left: The “full” DCM model that served as starting point for Bayesian model reduction. Black arrows represent intrinsic connections, coloured arrows denote driving inputs, and coloured dots represent modulatory inputs. Right: The resulting Bayesian Model Average thresholded at 99% parameter probability. Driving and between-region parameters are in units of Hz. Modulatory parameters in- or decrease connections in an additive manner. *: Significantly stronger modulation than other parameters (/Pp /> 0.99).

We found strong evidence for intrinsic connectivity within all three areas and pseudowords drove all three regions more strongly than words (Bayesian contrasts for TPC: *Pp*=0.999; IFG: *Pp*=1.0; vOTC: *Pp*=1.0). Crucially, effective connectivity from the left vOTC to the left TPC was significantly increased by effective TMS over the left TPC (modulation: 0.699; result: 0.464 Hz). This modulation was much stronger for effective than for sham TMS (modulation: 0.349; result: 0.113 Hz; Bayesian contrast between active and sham TMS: *Pp*=1.0). In other words, a disruption of the left TPC resulted in a stronger facilitatory drive from the left vOTC to the left TPC, which can be interpreted as a compensatory mechanism (see Discussion).

## 3. Discussion

### The role of the left TPC for pseudoword reading

In our present study, we explored whether the left TPC, a critical region for acquiring and establishing sound-letter mappings, is causally involved in pseudoword processing in adult readers. In other words, we tested if the typically reading brain still relies on this brain region when confronted with a task that requires access to singles sounds and syllables. Our findings show that a disruption of the left TPC impacts pseudoword processing. An inhibition of this area with TMS led to an increase of speech onsets for pseudowords. These behavioral changes were underpinned by a modulation of task-related activity and connectivity in the larger network for reading, likely reflecting compensatory reorganization in the reading network which may have helped to maintain processing. While the absence of a TMS-induced modulation of task-related activation in the stimulated area may be surprising, previous work demonstrated compensatory reorganization in distributed networks after inhibitory TMS for language and other cognitive tasks (see Hartwigsen & Volz, 2021 for review). Moreover, the behavioral relevance of remote changes induced by TMS in the language network has also been shown (Hartwigsen et al., 2017).

Although our findings support earlier TMS studies reporting NIBS-induced deteriorations or improvements in phonological performance (Turker &; Hartwigsen, 2021 a, b), our study is the first to provide direct evidence for a significant role of the left TPC for overt simple and complex pseudoword reading. The only comparable previous behavioral TMS study targeting the left TPC (Costanzo et al., 2012) reported an increase in reading accuracy but no differences in speech onsets. While facilitatory TMS effects on task performance are often interpreted as causal evidence for the contribution of the targeted area to a given task, such effects could also reflect the inhibition of task-irrelevant areas that compete for resources (Luber & Lisanby, 2014; see Bergmann & Hartwigsen, 2021 for discussion). The difference in the direction of results in the previous and present study is unclear. We believe that these differences could stem from a more precise neuronavigation in the present study, which may have resulted in targeting a different (sub-)region in our study, more complex stimuli (four instead of two or three syllables), a larger sample size and differences in stimulation parameters (continuous theta burst stimulation/cTBS instead of high-frequency rTMS). One other study tested pseudoword reading following TMS but in this language mapping study by Hauck and colleagues (2015a, b) several regions were targeted but no strong effect of TMS on the TPC could be found for pseudoword reading. Consequently, we here provide first evidence that the left TPC is causally involved in pseudoword processing since inhibitory TMS led to slower pseudoword reading and modulated communication within the reading network.

The observed TMS effect confirms neuroimaging findings indicating that the left TPC/inferior parietal cortex is still involved in spelling-sound conversion in adults (Taylor et al., 2013), despite evidence of this area’s contribution to various tasks including verbal working memory (Jonides et al., 1998) and executive processing (Binder et al., 2005). However, it also stands a bit in contrast to a meta-analysis on reading processing in child- and adulthood by Martin et al. (2015), who reported a strong engagement of the left TPC in children, but no significant activation in that area across neuroimaging studies with adults. Notably, the disruptive TMS effect of the present study was significant for pseudowords only (although words showed a trend towards delayed speech onsets). This observation may point to the relevance of the left TPC for phonological processing during reading. Alternatively, the stronger effect on pseudoword reading may have resulted from increased task difficulty for pseudoword relative to word reading. The behavioral results further showed a strong interaction between complexity and pseudowords, but this effect did not interact with TMS. Therefore, it is unlikely that the observed effect can selectively be explained by general task complexity.

Why would we see differences in speech onsets and not reading times? Even when asked to read written stimuli overtly, we process them before we start actual speech production, that is, as soon as we see them, we process and thus read them (we are unable to “not read” visual stimuli). Therefore, inhibiting an area that is crucial for access to letter-sound knowledge, will more likely impair this inner, ‘silent’ reading process than the task of speech production, which relies heavily on motor areas (Brown et al., 2005) and does not reflect reading mechanisms per se.

### Adaptive plasticity within the reading network

The present study is the first study that mapped TMS-induced behavioral changes on reading performance on the neural level with fMRI. Earlier research on language skills more generally suggests that TMS leads to plastic after-effects, such as large-scale changes on the network level affecting both local and remote activity within targeted networks, as well as interactions between other involved networks (Hartwigsen & Volz, 2021). As such, the perturbed brain can flexibly redistribute and functionally reorganize its computational capacities to compensate for the disruption of an area or the network. The present study adds first evidence for compensatory mechanisms within the typical reading network in terms of functional brain activation. We did not detect any significant TMS-related differences with univariate measures of brain activation, probably due to the observed strong interindividual variability in response to stimulation, which has already been reported for motor excitability in previous studies (see Hamada et al., 2013).

Nevertheless, we found TMS-induced changes in fine-grained multi-voxel activity patterns in the left IFC between effective and sham TMS, selectively for pseudoword (but not for word) reading. In terms of the functional role of the left IFC for reading, theories hold that it is crucial for phonological output resolution and rhyming during reading (Taylor et al., 2013; Brozdowski & Booth, 2021), as well as attention and working memory (Corbetta et al., 2002; Tops & Boksem, 2011). With pseudowords having a higher decoding demand and thus requiring more effort, specifically after inhibition of the left TPC, it is likely that differences in response patterns of the left IFC at least partially stem from higher demands on attention, executive functions, and cognitive control. Alternatively, the stronger contribution of left IFC to pseudoword reading after disruption of the left TPC might also reflect a shift in the balance towards another key node for reading, and thus reflect phonological processes per se. This explanation would be in line with previous TMS studies showing flexible redistribution between homologous areas (e.g., Hartwigsen et al., 2013; Jung & Lambon Ralph, 2016) or remote regions from the same specialized subnetwork (Hallam et al., 2016) during different language tasks (e.g., Hartwigsen, 2016). A stronger contribution of the left IFC after disruption of the left TPC likely reflects compensatory attempts in the network which helped to maintain task processing at a high level and may have prevented decreases in task accuracy.

Apart from the TMS-induced changes in response patterns, we found a shift in functional coupling between the left vOTC and the left TPC in response to inhibition of the latter. After effective TMS over the left TPC, as compared to sham stimulation, the left vOTC increased its facilitatory drive onto the left TPC. This is particularly interesting since the left vOTC plays a key role in orthographic processing and is vital for reading words and pseudowords (Jobard et al., 2003, Turker & Hartwigsen, 2021a). A functional engagement of this region for pseudoword processing could also be confirmed in the univariate analyses of this study. This observation of the left vOTC exerting a stronger influence on the left TPC during pseudoword processing could be interpreted as a compensatory mechanism. As such, functional connectivity between these two areas is most likely vital for successful and efficient decoding, so that a disruption of the left TPC requires an up-regulation of functional coupling to compensate for the increased demand posed by the task. This highlights the importance of considering within-network interactions when exploring TMS-induced effects on the neural level. When considering TMS-induced changes on task-related activity and connectivity, it is important to bear in mind that TMS is not “lesioning” an area and unlikely to completely “silence” processing in the targeted region. Consequently, a shift in the balance between different nodes in the respective network with a stronger contribution of another area may help to maintain processing at a relatively high level, despite the disruption (see Hartwigsen, 2018).

Overall, the present study emphasizes the relevance of the three core reading areas for pseudoword reading. It seems that the targeted region, the left TPC, is crucial for pseudoword processing, most likely for the processing of very complex stimuli. However, the reading network in typical adult readers is flexible enough to adapt to the disruption by increasing functional coupling between the left vOTC and the left TPC, and at the same time shifting functional brain activation in the left IFC. Since we still see a slight deterioration in pseudoword reading performance, as hypothesized, neural plasticity seems to be only partially successful at accommodating the induced disruption. This might still explain why the effects were not as large as expected, and only slightly affected response efficiency but not accuracy. The observed small behavioral effects most likely stem from large inter-individual variability in response to TMS. It seems that TMS responses are largely variable between individuals, which makes group-level analyses particularly challenging and explains both the small behavioral effects and the lack of univariate whole-brain activation differences in our study.

With respect to the contribution of our data to theoretical reading models, our study can be explained under the framework of two reading models, one being the connectionist framework of reading (Seidenberg et al., 2005), the other the dual-cascaded model of reading (DRC; Coltheart et al., 2001). The findings do not provide evidence for a hierarchical organization of the reading network, but they suggest a constant interaction between reading areas, more in line with connectionist accounts. Furthermore, the findings highlight that decoding recruits the left TPC, which is in line with earlier assumptions that unfamiliar word and pseudoword reading rely upon a dorsal reading stream, including recruitment of the vOTC, the left TPC and the left IFC (backed by structural connectivity research, e.g., Cummine et al., 2015). Since we did not observe any effects on word processing, this suggests that words might recruit a different route that does not require the left TPC. However, words could also be too automatized and robustly presented in the semantic lexicon as to respond to a TMS-induced disruption in the left TPC.

## 4. Conclusion

The present study provides first evidence that the left TPC is causally involved in overt pseudoword reading and confirms adaptive plasticity within the reading network. By combining rTMS with fMRI, we found that effective disruption led to (1) slower reading of pseudowords, manifested as a delay in speech onsets for simple and complex pseudowords, (2) a change in functional activation patterns in the left IFC as revealed by MVPA, and (3) an increase of functional coupling between the left vOTC and the left TPC. The latter two can be interpreted as compensatory mechanisms that show adaptive plasticity in the reading network in response to perturbation.

In summary, we report neurophysiological changes in response to TMS at the level of task-related activity and connectivity in addition to generally confirming earlier findings on causal contributions of the left TPC to reading-related phonological processes (e.g., Liederman et al., 2003; Costanzo et al., 2012). Even though we only used a single session intervention in this study, we could still see immediate effects on the behavioral and neural levels. The present findings can guide future studies and suggest new perspectives concerning the treatment of reading disorders, e.g., by designing multiple-session interventions for individuals with reading impairments. Overall, our study advances future experimental and translational applications of TMS in health and disease.

## 5. Methods

### 5.1. Participants

Participants were young, healthy, right-handed adults (N = 28; 13 females; range: 18-40 years, M_age_=25±4) with no prior history of psychiatric, neurological, hearing, or developmental disorders. All participants had nonverbal intelligence scores within the normal range or above (nonverbal IQ:≥ 91; CFT 20-R; Weiß, 2019). Sample size was determined based on comparable previous TMS studies (e.g., Kuhnke et al., 2020). Participants were either recruited via the participants database of the Max Planck Institute for Human Cognitive and Brain Sciences Leipzig (MPI CBS), or flyers, posters, and social media. Participation in all sessions was required for the respective participant’s data to be included in the study sample. Prior to participation, written informed consent was obtained from each subject. The study was performed according to the guidelines of the Declaration of Helsinki and approved by the local ethics committee of the University of Leipzig.

In accordance with the given governmental regulations and measures regarding the COVID-19 pandemic, participants were not allowed to have been to risk areas two weeks prior to study participation and were required to undergo a SARS-CoV-2 rapid antigen self-test upon arrival provided by the MPI CBS. Further, subjects were requested to sign a form stating that, in case of a positive result of the self-test, participation in the study had to be suspended.

### 5.2. Experimental procedure and behavioral reading assessment

The present study comprised one 3-hour behavioral testing session and two combined fMRI-TMS sessions (one for each TMS condition). During the behavioral testing session, we assessed nonverbal intelligence, working memory, and reading. However, for the present analysis, we only made sure that participants had normal nonverbal intelligence, reading and working memory. The *Culture Fair Test* (CFT 20-R; Weiß, 2019) was administered to test participants’ nonverbal intelligence. Verbal working memory was measured through digit span forward and digit span backward taken from the *Wechsler Adult Intelligence Scale* (WAIS-I; Petermann & Petermann, 2011). Additionally, nonword span (Mottier Test in the ZLT II-Zürcher Lesetest; Petermann & Daseking, 2019) was assessed. The test was terminated if the subject could not repeat 50% of syllables in one trial block. Silent text reading was assessed with the LGVT 5–12+ (Schneider et al., 2017), providing speed (number of words read), accuracy (ratio of filled gaps and correct items) and comprehension scores (number of correctly inserted words).

The TMS-fMRI sessions were separated by at least 7 days to prevent carry-over effects of TMS, and session order (sham or effective) was counterbalanced across participants. The study employed a 2×2×2 within-subject design with the factors TMS (effective stimulation, sham stimulation), stimulus type (words, pseudowords) and complexity (simple stimuli consisting of two syllables, complex stimuli consisting of four syllables) (for details of the experimental procedure, stimulation site and fMRI design, see **Figure 1**).

### 5.3. Transcranial magnetic stimulation (TMS)

To investigate the causal role of the left TPC for phonological processing, we applied “offline” (i.e, before the task) continuous theta burst stimulation (cTBS). cTBS applies bursts of 3 stimuli at 50 Hz repeated at intervals of 200 ms (5 Hz) for 40 seconds (total: 600 pulses) (Huang et al., 2005). Offline protocols can induce adaptive changes in brain activity and connectivity that outlast the stimulation for up to 60 minutes (Siebner & Rothwell, 2003). Participants underwent one effective and one sham (placebo) session. The sham condition mirrored the effective condition in terms of basic set-up and procedure, but a placebo coil (MCF-P-B65) was used, which features the same mechanical outline and acoustic noise as the effective coil but reduces the magnetic field strength by ∼ 80%.

Intensity of the stimulation was set at 90% of the individual resting motor threshold (rMT) The protocol for assessing the resting motor threshold was conducted in accordance with the standardized procedure proposed by Schutter and van Honk (2006). This procedure applies electromyography instead of purely visual observation of muscle twitch. rMT was determined as the lowest stimulation intensity producing at least 5 motor evoked potentials of ≥50 μV in the relaxed first dorsal interosseus muscle of the right hand when single-pulse TMS was applied over the hand region of left primary motor cortex 10 times.

The specific MNI coordinates for the left TPC (x=-49, y=-44, z=21) were calculated from three meta-analyses on reading impairments (Maisog et al., 2008; Richlan et al. 2009, 2011). To precisely target these coordinates in each individual participant, they were transformed from MNI to subject space using the *SPM12* software (Wellcome Trust Center for Neuroimaging, University College London, UK). We then used stereotactic neuronavigation (TMS Navigator, Localite GmbH, Sankt Augustin, Germany) to navigate the coil over the target area and maintain its location throughout stimulation. For neuronavigation, participants’ heads were co-registered onto their T1-weighted MR image before the stimulation sessions. T1 scans were obtained beforehand with a 3T MRI scanner (Siemens, Erlangen, Germany) using an MPRAGE sequence (176 slices in sagittal orientation; repetition time: 2.3 s; echo time: 2.98 ms; field of view: 256 mm; voxel size: 1 × 1 × 1 mm; no slice gap; flip angle: 9°; phase encoding direction: A/P).

### 5.4. Functional neuroimaging

#### Stimuli

We used an event-related mini-block design that used 400 stimuli altogether (200 simple and complex words; 200 simple and complex pseudowords). In each session, participants read randomly chosen 100 words (50 simple, 50 complex) and 100 pseudowords (50 simple, 50 complex) in mini-blocks (5 stimuli presented after another) aloud in the scanner (i.e., stimuli were not repeated in the second session to avoid remembering pseudowords). The 200 simple word stimuli consisted of two syllables and 4-6 letters and were taken from Schuster et al. (2015). As complex words, we chose the first 100 most frequent 4-syllabic words (10-14 letters) from the dlex database (http://www.dlexdb.de/). We excluded compound words and plurals but due to the small number of available complex words we had to include a few 3-syllabic words and plurals (in German the plural is usually constructed by adding a whole syllable). Pseudowords were then designed using Wuggy (http://crr.ugent.be/programs-data/wuggy) based on the simple and complex word lists. We excluded pseudowords that were too similar to real German words (<2 letters difference).

### Neuroimaging

Functional MRI data were collected on a 3T Siemens Magnetom Skyra scanner (Siemens, Erlangen, Germany) with a 32-channel head coil. Blood oxygenation level-dependent (BOLD) images were acquired with a single-echo BOLD EPI sequence (repetition time [TR]: 2s, echo time [TE]: 22ms; flip angle: 80°; field of view [FoV]: 204 mm; voxel size: 2.5 × 2.5 × 2.5 mm; bandwidth: 1794 Hz/Px; phase encoding direction: A/P; acceleration factor: 3). B0 field maps were acquired for susceptibility distortion correction using a spin-echo BOLD EPI sequence (TR: 8000ms; TE:50ms; flip angle: 90°; bandwidth: 1794 Hz/Px).

During fMRI, stimuli were presented for 2.5 seconds. We jittered the between-stimulus-interval, as well as the between-mini-block-interval. Each block lasted for 27.5 seconds and participants were in the scanner for around 25 minutes (see **Figure 1)**. Subjects were instructed to read out all stimuli as fast and correct as they could with as little head movement as possible. Subjects’ in-scanner responses were recorded and manually preprocessed with audacity. Speech on- and offsets were determined with Praat by four independent raters, two analyzing each audiofile in 50% of cases. We computed an interrater reliability >0.85 suggesting that determination of speech on- and offsets was very similar between raters. For the following analyses, we averaged speech onsets across raters if they were rated by more than one person. Accuracy for all trials was checked by a third person.

### 5.5. Data analysis

#### 5.5.1. Linear Mixed Model

Speech onsets and response accuracy of each trial were analysed with generalised linear mixed models (GLMM) using glmmTMB 1.1.2.3 (Brooks et al., 2017) in R 4.0.5. To circumvent the need to transform reaction times to satisfy normality assumptions, reading times were modelled using a Gamma distribution with the identity link function (Lo & Andrews, 2015). The accuracy of each trial (correct versus incorrect) was modelled as a binary response using a binomial distribution and logit link function. All models included as fixed effects the three-way interaction between TMS (sham/active), Pseudoword (pseudoword/word), and Complexity (simple/complex), and all lower order terms. A maximal random effects structure was used for all models with subject as the grouping variable to avoid inflated Type I errors (Barr et al., 2013). The resulting GLMM for speech onsets included random intercepts, and random slopes for the interaction between TMS, Complexity and Pseudoword, as well as all lower order terms. Likewise, random intercepts and random slopes for TMS and Complexity were included in the GLMM for response accuracy. The significance of each variable was assessed using the Wald test, and marginal effects were calculated using a step-down simple effects analysis.

#### 5.5.2. fMRI analysis

##### Preprocessing

MRI preprocessing was performed using *fMRIprep* (version 20.2.1; Esteban et al. 2019). Anatomical T1-weighted images were corrected for intensity non-uniformity (using *N4BiasFieldCorrection* from ANTs 2.3.3), skull-stripped (using *antsBrainExtraction* from ANTs 2.3.3), segmented into gray matter, white matter and cerebrospinal fluid (using *fast* in FSL 5.0.9), and normalized to MNI space (MNI152NLin2009cAsym; using *antsRegistration* in ANTs 2.3.3). Brain surfaces were reconstructed using *reconall* (FreeSurfer 6.0.1).

Functional BOLD images were co-registered to the anatomical image (using *bbregister* in FreeSurfer 6.0.1), distortion corrected based on B0-fieldmaps (using *3dQwarp* in AFNI 20160207), slice-timing corrected (using *3dTshift* from AFNI 20160207), motion corrected (using *mcflirt* from FSL 5.0.9), normalized to MNI space (via the anatomical-to-MNI transformation), and smoothed with a 5 mm^3^ FWHM Gaussian kernel (using *SPM12*; Wellcome Trust Centre for Neuroimaging; http://www.fil.ion.ucl.ac.uk/spm/). Moreover, physiological noise regressors were extracted using the anatomical version of *CompCor* (aCompCor, Behzadi et al., 2007).

##### Univariate analyses

We performed a whole-brain random-effects group analysis based on the general linear model (GLM), using the two-level approach in *SPM12*. At the first level, individual participant data were modeled separately. The participant-level GLM included regressors for the 4 experimental conditions, modelling trials as box car functions (2.5 s duration) convolved with the canonical HRF. Only correct trials were analyzed, error trials were modeled in a separate regressor-of-no-interest. Nuisance regressors included 24 motion regressors (the 6 base motion parameters + 6 temporal derivatives of the motion parameters + 12 quadratic terms of the motion parameters and their temporal derivatives), individual regressors for time points with strong volume-to-volume movement (framewise displacement > 0.9; Siegel et al. 2014), and the top 10 aCompCor regressors explaining the most variance in physiological noise. The data were subjected to an AR(1) auto-correlation model to account for temporal auto-correlations, and high-pass filtered (cutoff 128 s) to remove low-frequency noise.

Contrast images for each participant were computed at the first level. At the second level, these contrast images were submitted to one-sample or paired t-tests (to test for interactions). For all second-level analyses, a gray matter mask was applied, restricting statistical tests to voxels with a gray matter probability > 0.1 (MNI152NLin2009cAsym gray matter template in *fMRIprep*). All activation maps were thresholded at a voxel-wise p < 0.001 and a cluster-wise p < 0.05 FWE-corrected.

##### Multivariate pattern analysis (MVPA)

As univariate analyses are insensitive to information represented in fine-grained, multi-voxel activation patterns (Haxby et al., 2014), we additionally performed a multivariate pattern analysis (MVPA) using *The Decoding Toolbox* (Hebart et al., 2015) implemented in *Matlab* (version 2021a). Our MVPA aimed to test whether effective TMS over TPC, as compared to sham TMS, modulated activity patterns in the stimulated or other, remote brain regions. We employed searchlight MVPA, moving a spherical region-of-interest (or “searchlight”) of 5 mm radius through the entire brain (Kriegeskorte et al., 2006). At each searchlight location, a machine-learning classifier (an L2-norm support vector machine; C=1) aimed to decode between effective and sham TMS, separately for words and pseudowords. We used leave-one-*participant*-out cross validation (CV), training on the activation patterns from n-1 participants and testing on the left-out participant (yielding 28 CV-folds). For statistical inference, we performed a permutation test across the accuracy-minus-chance maps of the different CV-folds (using *SnPM13*; as proposed by Wang et al., 2021), thresholded at a voxel-wise p < 0.001 and a cluster-wise p < 0.05 FWE-corrected (as in our univariate analyses). Activity patterns comprised beta estimates for each mini-block of every participant.

##### Dynamic Causal Modelling (DCM)

Finally, we performed dynamic causal modelling (DCM; Friston et al., 2003) to investigate TMS-induced changes in effective connectivity (i.e., directed causal influences) between the core nodes of the reading network. DCM estimates a model of effective connectivity between brain regions to predict a neuroimaging time series. A DCM consists of three types of parameters: 1) “intrinsic” (i.e., condition-independent) directed connections between brain regions, 2) “modulatory inputs” that change connection strengths during a certain experimental manipulation, and 3) “driving inputs” that drive activity in the network. The goal of DCM is to optimize a tradeoff between model fit (of the predicted to observed time series) and complexity (i.e., deviation of model parameters from their prior expectations), measured by the model evidence (Kahan & Foltynie, 2013; Zeidman et al., 2019a).

We performed a two-level analysis using Parametric Empirical Bayes (PEB) and Bayesian Model Reduction (BMR)—the current “standard practice for group DCM studies” (Friston et al., 2016). At the first level, a “full model” was specified and estimated for each participant (see Results section). Regions included in the model were the left TPC (the stimulated region), left vOTC, and left IFG. The three regions were defined functionally in each individual participant as the top 10% most activated voxels for [all sham trials > rest] within 20 mm spheres around the MNI peak coordinates in a meta-analysis of reading in adults (Martin et al., 2015): left TPC = -49-44 21; left IFG = -52 20 18; left vOTC = -42 -68 -22. All regions were restricted to the cerebral gray matter. The first eigenvariate of the BOLD time series of each region was extracted and adjusted for effects-of-interest (all experimental conditions) using our participant-level GLM (see Univariate analyses). DCM inputs were mean-centered, so that the intrinsic connections reflected the mean connectivity across experimental conditions (Zeidman et al., 2019a).

At the second level, DCM parameters of individual participants were entered into a GLM— the PEB model—that decomposed interindividual variability in connection strengths into group effects and random effects (Zeidman et al., 2019b). BMR then compared the full model against numerous reduced models that had certain parameters “switched off” (i.e., prior mean and variance set to 0) (Friston et al., 2016). Finally, we computed the Bayesian model average (BMA), the average of parameter values across models weighted by each model’s posterior probability (Pp) (Penny et al., 2007). This approach is preferred over exclusively assessing the parameters of the “best” model as it accommodates uncertainty about the true underlying model (Friston et al., 2016; Dijkstra et al., 2017). The BMA was thresholded to only retain parameters with a Pp > 99% (cf. Zeidman et al., 2019b; Kuhnke et al., 2021). For each modulatory input, we calculated the resulting connectivity value (in Hz) using formula 3 in Zeidman et al. (2019a). Finally, to determine whether one experimental condition modulated a certain connection more strongly than another, we directly compared different parameters on the same connection using Bayesian contrasts (Dijkstra et al., 2017; Kuhnke et al., 2021).

## 6. Acknowledgements

The present work was supported by the Max Planck Society and the Humboldt Foundation.

